# Widespread Inter-Chromosomal Epistasis Regulates Glucose Homeostasis and Gene Expression

**DOI:** 10.1101/132175

**Authors:** Anlu Chen, Yang Liu, Scott M. Williams, Nathan Morris, David A. Buchner

## Abstract

The relative contributions of additive versus non-additive interactions in the regulation of complex traits remains controversial. This may be in part because large-scale epistasis has traditionally been difficult to detect in complex, multi-cellular organisms. We hypothesized that it would be easier to detect interactions using mouse chromosome substitution strains that simultaneously incorporate allelic variation in many genes on a controlled genetic background. Analyzing metabolic traits and gene expression levels in the offspring of a series of crosses between mouse chromosome substitution strains demonstrated that inter-chromosomal epistasis was a dominant feature of these complex traits. Epistasis typically accounted for a larger proportion of the heritable effects than those due solely to additive effects. These epistatic interactions typically resulted in trait values returning to the levels of the parental CSS host strain. Due to the large epistatic effects, analyses that did not account for interactions consistently underestimated the true effect sizes due to allelic variation or failed to detect the loci controlling trait variation. These studies demonstrate that epistatic interactions are a common feature of complex traits and thus identifying these interactions is key to understanding their genetic regulation.

---

The genetic basis of complex traits and diseases results from the combined action of many genetic variants [1]. However, it remains unclear whether these variants act individually in an additive manner or via non-additive epistatic interactions. Epistasis has been widely observed in model organisms such as *S. cerevisiae* [2,3], *C. elegans* [4], *D. melanogaster* [5] and *M. musculus* [6]. However, it has been more difficult to detect in humans, potentially due to their diverse genetic backgrounds, low allele frequencies, limited sample sizes, complexity of interactions, insufficient effect sizes, and methodological limitations [7,8]. Nonetheless, a number of genome-wide interaction-based association studies in humans have provided evidence for epistasis in a variety of complex traits and diseases [9–15]. However, concerns remain over whether observed epistatic interactions are due to statistical or experimental artifacts [16,17].

To better understand the contribution of epistasis to complex traits, we studied mouse chromosome substitution strains (CSSs) [18]. In CSSs, a single chromosome in a host strain is replaced by the corresponding chromosome from a donor strain. This provides an efficient model for mapping quantitative trait loci (QTLs) on a fixed genetic background. This is in contrast to populations with many segregating variants such as advanced intercross lines [19], heterogeneous stocks [20], or typical analyses in humans. Given the putative importance of genetic background effects in complex traits [21,22], we hypothesized the fixed genetic backgrounds of CSSs can provide a novel means for detecting genetic interactions on a large-scale [18,23]. Previous studies of CSSs with only a single substituted chromosome suggested that non-additive epistatic interactions between loci were a dominant feature of complex traits [6]. However, to identify the interacting loci, or at least their chromosomal locations, requires the analysis of genetic variation in multiple genomic contexts [24]. We thus extended the analysis of single chromosome substitutions by analyzing a series of CSSs with either one or two substituted chromosomes, collectively representing the pairwise interactions between genetic variants on the substituted chromosomes. This experimental design can directly identify and map loci that are regulated by epistasis by analyzing the phenotypic effects of genetic variants on multiple fixed genetic backgrounds. Here we report the widespread effects of epistasis in controlling complex traits and gene expression. The detection of true epistatic interactions will improve our understanding of trait heritability and genetic architecture as well as providing insights into the biological pathways that underlie disease pathophysiology [25]. Knowing about epistasis will also be essential for guiding precision medicine-based decisions by interpreting specific variants in appropriate contexts.

## Results

### Contribution of epistasis to metabolic traits

Body weight and fasting plasma glucose levels were measured in a total of 766 control and CSS mice (S1, S2 Tables, S1 Fig.). The CSSs included 240 mice that were heterozygous for one A/J-derived chromosome and 444 mice that were heterozygous for two different A/J-derived chromosomes, both on otherwise B6 backgrounds. The CSSs with two A/J-derived chromosomes represented all pairwise interactions between the individual A/J-derived chromosomes. For example, comparisons were made between strain B6, strains (B6.A3 × B6)F1 and (B6 × B6.A10)F1 which were both heterozygous for a single A/J-derived chromosome (Chr. 3 and 10, respectively), and strain (B6.A3 × B6.A10)F1 which was heterozygous for A/J-derived chromosomes 3 and 10 (S2 Fig.). A complete list of the strains analyzed is shown in Table S2. Quantitative trait loci (QTLs) were identified for both body weight and plasma glucose levels that were due to main effects and interaction effects. Due to the study design, only QTLs with dominant effects could be assessed.

Omnibus tests for main effects on body weight indicated that some of the chromosome substitutions individually influenced body weight (males p=0.0028; females p=0.0008; meta p=1.4e-05). Similarly, omnibus tests for main effects on plasma glucose levels demonstrated a significant effect of the chromosome substitutions (males p=0.0082; females p=0.00011; meta p=1.4e-05). QTLs with main effects on body weight were mapped to chromosomes 8 (Main Effect: 1.23g; Average Effect: 1.02g) and 17 (Main Effect: -1.13g; Average Effect: (-1.11g) (S3 Table). Note that we define main effects as the effect of a chromosome substitution as estimated by a model which includes all pairwise interaction terms, thus taking into account context-dependent genetic background effects. In contrast, the average effect is estimated using a model that does not include any interaction terms; the latter is similar to the analyses performed in a typical GWAS study. QTLs with main effects on fasting glucose were mapped to chromosomes 3 (Main Effect: 25.0 mg/dL; Average Effect: 9.61 mg/dL), 5 (Main Effect: 15.6 mg/dL; Average Effect: 6.02 mg/dL), and 4 (Main Effect: 17.5 mg/dL; Average Effect: 6.61 mg/dL) (S3 Table).

Omnibus tests for interaction effects on body weight were not significant (males p= 0.19; females p= 0.83; meta p= 0.44), and therefore epistatic interactions on body weight were not further investigated. However, omnibus tests for interaction effects on plasma glucose demonstrated the importance of epistasis in regulating this trait (males p= 0.002; females p= 0.003; meta p= 8.99e-05). In fact, among the males and females respectively, epistasis accounted for 43% (95% confidence interval: 23%-75%) and 72% (95% confidence interval: 37%-97%) of the heritable effects on plasma glucose levels. The discrepant results for the contribution of interactions to body weight and plasma glucose are likely reflected in the difference between whether QTLs for these traits were detected using the main effect model or the average effect model (S3 Table). For plasma glucose, only 1 of the 3 QTLs identified using the main effect model was also identified using the average effect model, and no new QTLs were identified with the average effect model. In contrast, both of the QTLs for body weight identified using the main effect model were also identified using the average effect model, and 2 new QTLs were identified on chromosomes 6 and 10. This suggests that for a trait regulated by epistatic interactions, the ability to successfully identify QTLs is greatly enhanced by accounting for these interactions. However, for a trait regulated primarily by additive effects, a model incorporating interactions can be detrimental to QTL identification.

To identify specific epistatic interactions, we tested explicit hypotheses for inter-chromosomal pairwise interactions on plasma glucose levels. Among the 15 CSS crosses analyzed, 5 crosses demonstrated inter-chromosomal epistatic interactions that altered plasma glucose levels (Fig. 1, S3, S4 Figs.). Interestingly, in all 5 crosses demonstrating interactions, one chromosome substitution increased fasting glucose levels relative to the control B6 strain. These main effects raised plasma glucose levels by an average of 12.3 mg/dL in males and 17.8 mg/dL in females. However, in all 5 observed interactions the average plasma glucose levels in the double CSSs were closer to the control B6 strain than any single CSS was. Furthermore, in 4 of the 5 interactions, the plasma glucose levels in the double CSS did not differ statistically from the control strain B6 (p value > 0.1). Thus, the chromosome substitution driving the increase in plasma glucose on a B6 background had no effect on glucose levels when the genetic background was altered by the second chromosome substitution.

**Figure 1.**
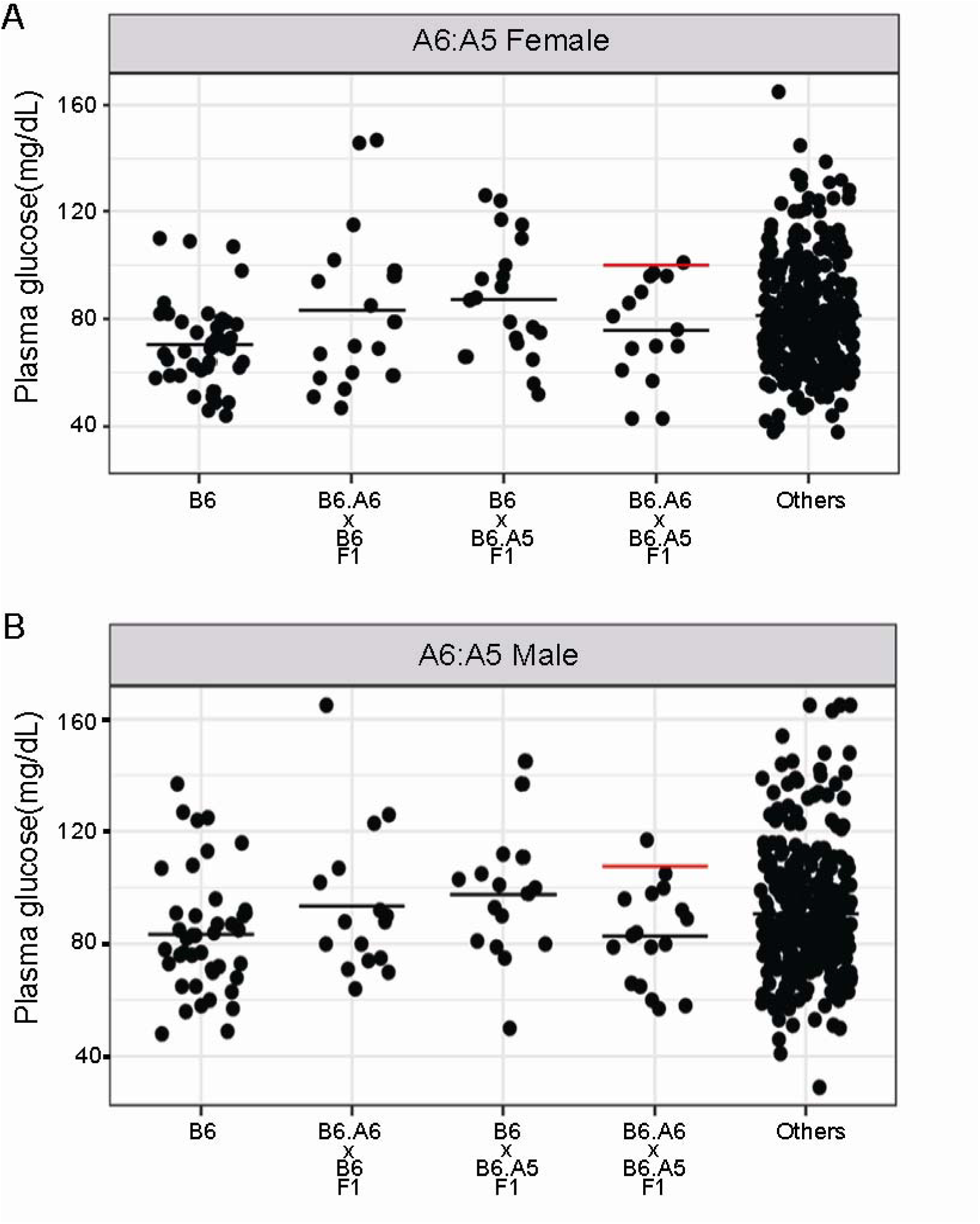
Inter-chromosomal epistasis between chromosomes 5 and 6 regulates fasting plasma glucose levels in mice. Plasma glucose levels were measured in 5-week-old mice that were fasted overnight. Each dot represents the glucose level of a single mouse. “Others” represents the data from all mice in this study excluding the other 4 strains shown in that panel. The black horizontal line indicates the mean glucose level for each group. The red horizontal line indicates the predicted trait level based on a model of additivity.

### Regulation of gene expression by epistasis

As hepatic gluconeogenesis is a key determinant of plasma glucose levels in healthy insulin-sensitive mice [26], the hepatic gene expression patterns of control and CSS mice were analyzed to better understand the molecular mechanisms underlying the epistatic regulation of plasma glucose. The RNA-Seq data was filtered for genes expressed in the liver, leaving 13,289 genes that were tested for differential expression associated with both main and interaction effects. A total of 6,101 main effect expression QTLs (meQTLs) were identified (FDR < 0.05) (Fig. 2, S4 Table). Those meQTL genes located on the substituted chromosome were classified as cis-meQTLs (Fig. 2, red) whereas the meQTL genes not located on the substituted chromosome were classified as trans-meQTLs (Fig. 2, blue). Among all possible genes regulated by a cis-meQTL, on average 11.48% of these genes in each strain had a cis-meQTL (range: 5.54% - 22.09%) (S5 Table). Similarly, among all possible genes regulated by a trans-meQTL, on average 5.42% (range: 0.08% to 19.26%) of these genes were regulated by a trans-meQTL (S5 Table). The percentage of cis- and trans-meQTLs in each strain demonstrated a strong positive linear correlation (Spearman’s r = 1.0) but the proportion of cis-eQTLs was always greater than the proportion of trans-eQTLs. Strain (B6 × B6.A8)F1 had both the highest percentage of genes with cis-meQTLs (22.09%) and trans-meQTLs (19.26%), whereas strain (B6 × B6.A5)F1 had both the lowest percentage of genes with cis-meQTLs (5.54%) and trans-meQTLs (0.08%). This suggests that trans-meQTLs are being driven by the cumulative action of many cis-effects rather than a single or small number or major transcriptional regulators (S5 Fig.). Among the genes regulated by a meQTL(s), 41.98% (1615 out of 3847) were regulated by multiple meQTLs (Range: 2-6) (S6, S7 Tables). For example, *Brca2* is regulated by 5 trans-meQTLs mapped to chromosomes 4, 6, 8, 10 and 14 (S6 Fig., S7 Table), demonstrating that hepatic *Brca2* expression is regulated by allelic variation throughout the genome. In addition to the well-known role of *Brca2* in breast cancer susceptibility, *Brca2* has been implicated in hepatocellular carcinoma risk [27–29].

**Figure 2.**
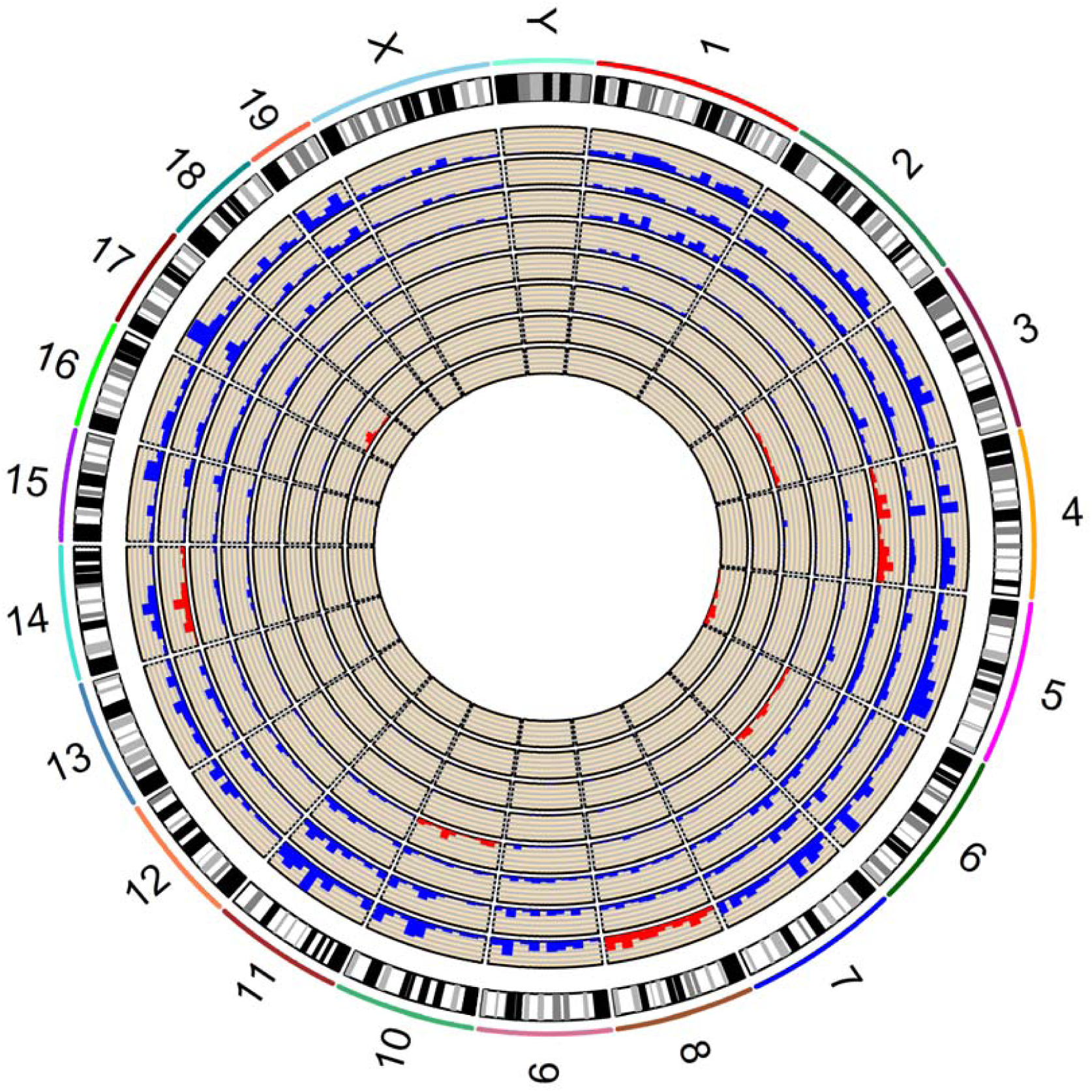
Identification of meQTLs that regulate hepatic gene expression. A circos plot of meQTL locations in the genome where each layer of the circle represents the comparison between a CSS strain and control B6 mice. From the inner circle, the CSS strains are (B6 × B6.A5)F1, (B6.17 × B6)F1, (B6.A3 × B6)F1, (B6.A6 × B6)F1, (B6 × B6.A10)F1, (B6 × B6.A4)F1, (B6.A14 × B6)F1 and (B6 × B6.A8)F1. Cis-meQTLs and trans-meQTLs are marked with red and blue, respectively. The width of each chromosome is proportional to its physical size. The height of each meQTL bar is proportional to the number of meQTLs in that genomic interval.

In addition to the meQTLs regulated by substitution of a single chromosome, the analysis of double CSSs enabled the detection of eQTLs with additive and interaction effects between the substituted chromosomes. The expression of *Zkscan3* represents an example of additivity, with the substitution of A/J-derived chromosomes 8 and 17 each individually increasing the expression of *Zkscan3* relative to control B6 mice (S7 Fig.). In the double CSS strain (B6.A17 × B6.A8)F1, the effects of each individual chromosome substitution are combined in an additive manner to result in yet higher expression than either of the single CSSs (S7 Fig.). The additive effects of the *Zkscan3* meQTLs detected by RNA-Seq were confirmed by quantitative reverse transcription PCR (S7 Fig.), as were 4/5 additional meQTLs demonstrating additivity (S8 Fig.).

In addition to examples of additivity, interaction expression QTLs (ieQTLs) were identified that were jointly regulated by genetic variation on two substituted chromosomes. The ieQTLs, similar to the meQTLs, were divided into cis-ieQTLS and trans-ieQTLs, with cis-ieQTLs defined by differentially expressed genes located on either one of the two substituted chromosomes and trans-ieQTLs representing differentially expressed genes that are not located on either substituted chromosome. A total of 4,283 ieQTLs were identified (S9 Table). Among all possible genes regulated by a cis-ieQTL or trans-ieQTL, 2.01% and 2.16% of genes were regulated by a cis- or trans-ieQTL respectively (Table 1). The combination of A/J-derived chromosomes 8 and 14 yielded the most ieQTLs (n=2,305) including cis-ieQTLs regulating the expression of 17.56% of all genes on chromosomes 8 or 14 and trans-ieQTLs regulating the expression of 17.32% of all genes throughout the remainder of the genome. Overall, the ieQTLs demonstrated a similar positive linear correlation as the meQTLs (Spearman’s r = 0.92) (S8 Fig.), although there was no enrichment for cis-ieQTLs. Among the genes regulated by an ieQTL(s), 32.35% (945 out of 2921) were regulated by multiple ieQTLs (Range: 2-7) (S10, S11 Tables). For example, *Agt* expression is decreased in strain (B6.A8 × B6)F1 relative to control B6 mice; however, interactions between alleles on chromosome 8 and chromosomes 6, 3, 17, and 14 all result in expression levels of *Agt* that did not differ from the control strain (Fig. 3).

**Table 1.** Interection effects on gene expression

| Cross | Cis-ieQTLs |  |  | Trans-ieQTLs |  |  | Subtypes of ieQTLs |  |  |  |
| --- | --- | --- | --- | --- | --- | --- | --- | --- | --- | --- |
|  | Percentage of genes with cis-ieQTL | Number of cis-ieQTLs | Number of genes on substituted chromosome | Percentage of genes with trans-ieQTL | Number of trans-ieQTLs | Number of genes on other chromosome | Percentage of genes with synergistic ieQTLs | Number of synergistic ieQTLs | Percentage of genes with antagonistic ieQTLs | Number of antagonistic ieQTLs |
| A14:A8 | 17.56% | 199 | 1133 | 17.32% | 2106 | 12156 | 6% | 129 | 94% | 2176 |
| A6:A8 | 6.81% | 96 | 1409 | 5.89% | 700 | 11880 | 2% | 15 | 98% | 781 |
| A14:A4 | 3.86% | 48 | 1244 | 3.57% | 430 | 12045 | 6% | 31 | 94% | 447 |
| A6:A10 | 1.81% | 24 | 1325 | 0.58% | 69 | 11964 | 1% | 1 | 99% | 92 |
| A14:A10 | 1.62% | 17 | 1049 | 2.05% | 251 | 12240 | 1% | 3 | 99% | 265 |
| A6:A4 | 0.86% | 13 | 1520 | 0.58% | 68 | 11769 | 0% | 0 | 100% | 81 |
| A3:A8 | 0.77% | 11 | 1434 | 0.77% | 91 | 11855 | 2% | 2 | 98% | 100 |
| A3:A10 | 0.44% | 6 | 1350 | 0.56% | 67 | 11939 | 0% | 0 | 100% | 73 |
| A17:A8 | 0.23% | 3 | 1329 | 0.00% | 0 | 11960 | 0% | 0 | 100% | 3 |
| A17:A4 | 0.14% | 2 | 1440 | 0.48% | 57 | 11849 | 2% | 1 | 98% | 58 |
| A17:A10 | 0.08% | 1 | 1245 | 0.03% | 4 | 12044 | 0% | 0 | 100% | 5 |
| A14:A5 | 0.00% | 0 | 1410 | 0.17% | 20 | 11879 | 0% | 0 | 0% | 0 |
| A17:A5 | 0.00% | 0 | 1606 | 0.00% | 0 | 11683 | 0% | 0 | 0% | 0 |
| A3:A5 | 0.00% | 0 | 1711 | 0.00% | 0 | 11578 | 0% | 0 | 0% | 0 |
| A6:A5 | 0.00% | 0 | 1686 | 0.00% | 0 | 11603 | 0% | 0 | 100% | 20 |
| All | 2.01% | 420 | 20891 | 2.16% | 3863 | 178444 | 4% | 182 | 96% | 4101 |

**Figure 3.**
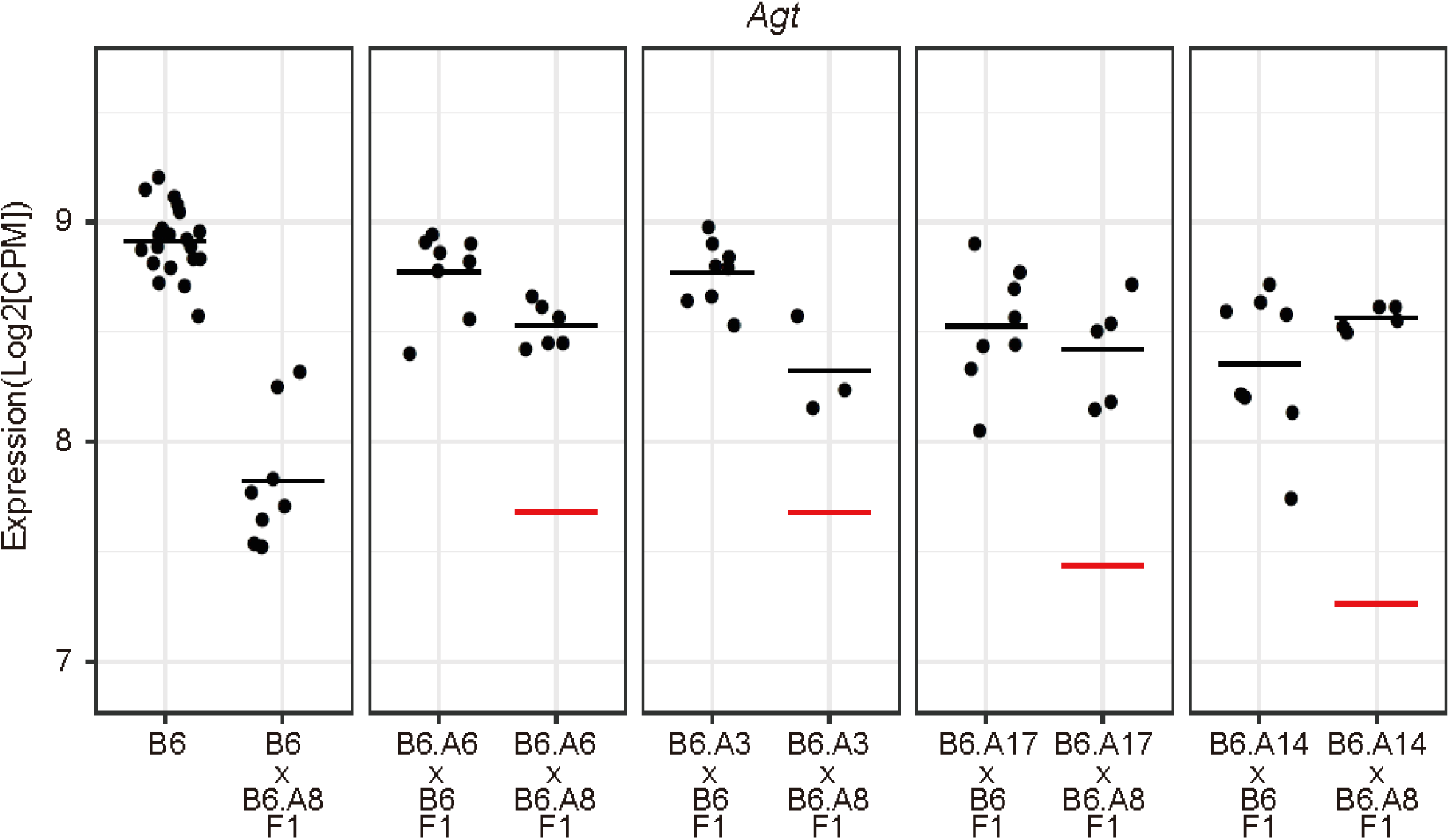
Identification of 4 ieQTLs that regulate the hepatic expression of Agt. Gene expression levels of Agt in the liver are shown for strain B6, 5 single CSS strains, and 4 double CSS strains. Each dot represents Agt expression in an individual mouse. The mean value for each strain is indicated by a solid line. The expected expression level of Agt in the double CSS strains based on a model of additivity is indicated with a red line. The Agt gene is located on mouse chromosome 8.

### Context-dependent effects on gene expression

We next tested whether the interaction effects on gene expression were synergistic (positive epistasis) or antagonistic (negative epistasis) (S9 Fig.). Synergistic refers to an increased difference in gene expression levels between the double CSS and the control B6 strain beyond that expected based on an additive model, whereas antagonistic refers to a decreased difference. The regulation of *Agxt* was an example of an antagonistic interaction, with main effects from substituted chromosomes 6 and 8 each individually decreasing *Agxt* expression, whereas this effect was lost in the double chromosome substitution strain (Fig. 4A). In contrast, the regulation of *Cyp3a16* represented an example of synergistic interaction with the detection of an ieQTL in the absence of a meQTLs (Fig. 4B). Among the ieQTLs, antagonistic interactions accounted for 96% (n=4101) while synergistic interactions accounted for 4% (n=182) (Table 1). Remarkably, for 80% of the antagonistic interactions (3285/4101), gene expression in one or both of the single CSSs differed from the control B6 strain (a meQTL), whereas expression in the double CSS reverted to control levels (p > 0.1 relative to strain B6). To again validate the RNA-Seq data using an independent method, RT-qPCR was performed for a subset of genes with antagonistic (n=13) and synergistic (n=10) interactions. Replication by RT-qPCR confirmed the detection of epistasis in 61% (p <0.05) of the genes tested (Antagonistic: 8/13; Synergistic: 6/10) (S8 Table).

**Figure 4.**
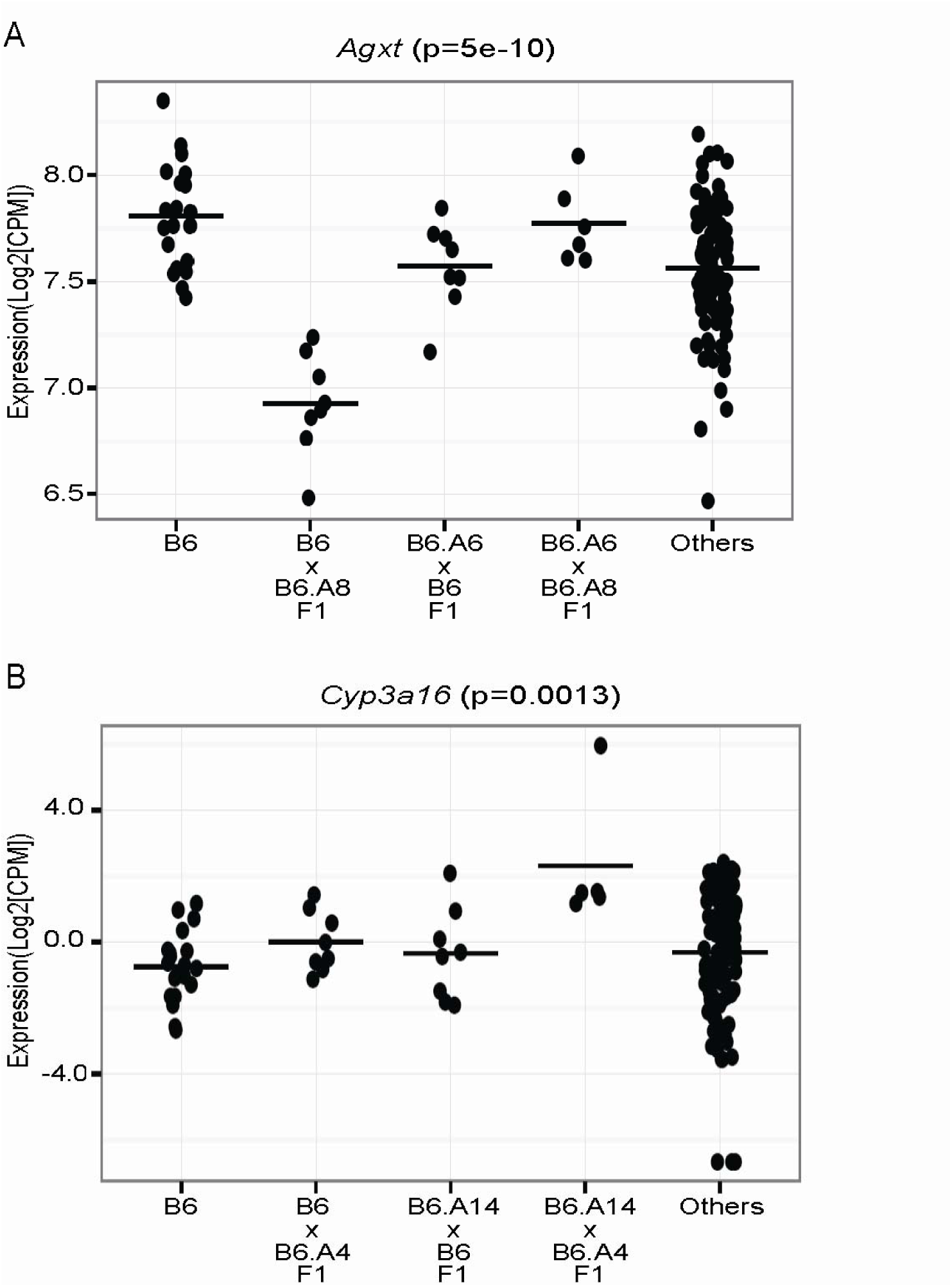
Examples of synergistic and antagonistic ieQTLs. Each dot represents the gene expression data from one mouse. The horizontal bar indicates the mean value for each strain (A) An antagonistic ieQTL regulates the expression of *Agxt* in the liver. (B) A synergistic ieQTL regulates the expression of *Cyp3a16* in the liver.

### Significant contribution of epistasis to trait heritability

Given that the ieQTLs regulated approximately 2% of all genes expressed in the liver (Table 1), we sought to quantify the contribution of genetic interactions to the heritable component of all genes. First, an omnibus test identified 6,684 genes out of the 12,325 genes expressed in the liver for which there was evidence of genetic control within the population of CSSs. The average proportion of heritable variation attributable to interactions across these genes was 0.56 (1^st^ quartile: 0.43 – 3^rd^ quartile: 0.68) (Fig. 5A). When the same analysis was restricted to only genes with a statistically significant contribution of interactions to gene expression levels (n=3,236 genes), the proportion of heritable variation attributable to interactions increased to 0.66 (1^st^ quartile: 0.56 – 3^rd^ quartile: 0.74) (Fig. 5B).

**Figure 5.**
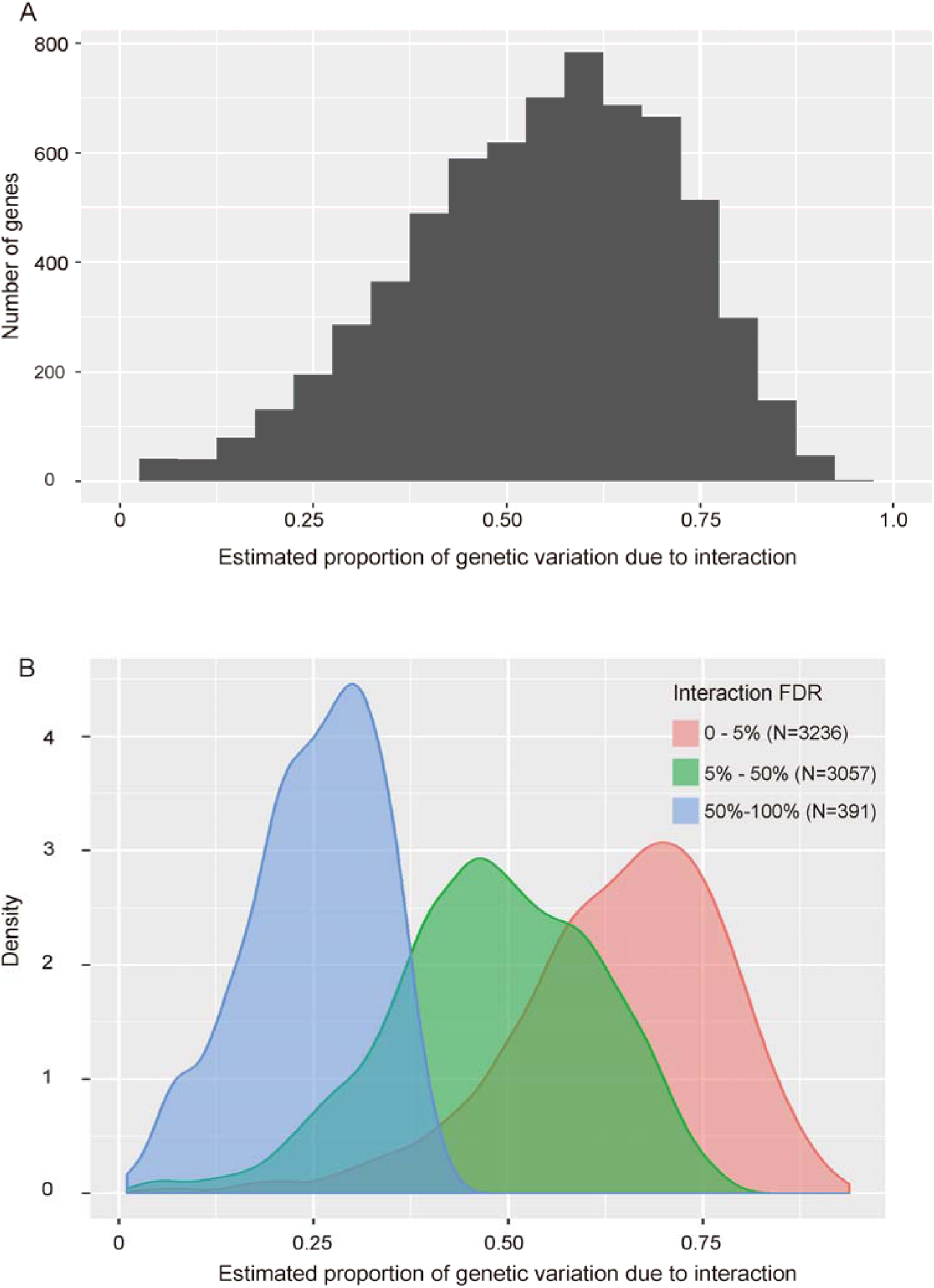
Contribution of epistasis to the genetic regulation of hepatic gene expression. (A) Diagrams representing the estimated proportion of genetic variation due to interactions for (A) all genes expressed in the mouse liver whose expression was under genetic control in the CSS strains studied and (B) the same data segregated based on the statistical evidence supporting an effect of interaction on gene expression.

## Discussion

CSSs, which have a simplified and fixed genetic background, were used to identify widespread and likely concurrent epistatic interactions. This systematic analysis of mammalian double CSSs demonstrated that epistatic interactions controlled the majority of the heritable variation in both fasting plasma glucose levels and hepatic gene expression (Fig. 5). Among genes expressed in the liver, the expression level of 24% were regulated, at least in part, by epistasis (Fig. 5). This number is remarkable considering that only dominant effects were tested, only a single tissue and time point were examined, allelic variation from only two inbred strains of mice were included, and only 15 pairwise strain combinations of CSSs were tested out of a possible 462 combinations of double CSSs. The prevalence of epistatic interactions provides a potential molecular mechanism underlying the highly dependent nature of complex traits on genetic background [21,22,30,31]. Interpreting the effect of individual allelic variants will thus be severely limited by population-style analyses that fail to account for possible contextual effects. Nonetheless, progress is being made in this field, including in diseases such as multiple sclerosis (MS), which is a complex genetic disease whose risk is highly associated with family history [32]. For example, MS risk alleles in DDX39B (rs2523506) and IL7R (rs2523506A) together significantly increase MS risk considerably more than either variant independently [15]. Based on the considerable number of interactions detected in the CSS crosses, context-dependent interactions such as that between DDX39B and IL7R in MS are likely widespread and may therefore represent a significant source of missing heritability for complex traits and diseases [33,34].

Although epistasis was a dominant factor regulating fasting glucose levels, the same effect was not detected in the regulation of body weight. It is not clear if this is due to different genetic architectures between these two traits or whether this was due to the limited genetic variation between the B6 and A/J strains. The body weight studies were conducted in mice fed a standard rodent chow, whereas differences in body weight between strains B6 and A/J are significantly more pronounced when challenged with a high-fat diet [35,36]. Alternately, a recent meta-analysis of trait heritability in twin studies identified significant variation in the role of additive and non-additive variation among different traits, with suggestive evidence for non-additive effects in 31% of traits [37]. Among the traits analyzed, genetic regulation of neurological, cardiovascular, and ophthalmological traits were among the most consistent with solely additive effects, whereas traits related to reproduction and dermatology were more often consistent with non-additive interactions. Among the metabolic traits studied, 40% of the 464 traits studied were consistent with a contribution of non-additive interactions [37]. It is interesting to speculate whether some traits that may have a more direct effect on fitness (e.g. reproduction) are more likely to involve multiple non-additive effectors in order to maintain a narrow phenotypic or developmental range [38].

Although many inter-chromosomal non-additive interactions were identified in mice, it remains unclear whether these interactions are attributable to bigenic gene-gene interactions or to higher-order epistasis involving multiple loci located on a substituted chromosome. Studies in yeast that dissected the genetic architecture of epistasis demonstrated that gene-gene interactions played a minor role among the heritable effects attributable to epistasis, thus primarily implicating higher order interactions [2]. Yet, other studies in yeast that methodically tested pairs of gene knockouts for interactions identified a number of gene-gene interactions [39]. Additional evidence for both high-order epistasis with three, four, and even more mutations [40] as well as bigenic gene-gene interactions [41] have been identified and it seems likely that both will underlie interactions detected in the CSS studies. However, to formally test this and determine the relative contribution of each, higher resolution genetic mapping of the epistatic interactions will be necessary to better understand their molecular nature [42].

Perhaps the most significant outcome of the epistasis detected was the high degree of constancy in the light of context dependence, such that the interactions usually returned trait values to the levels detected in control mice. Remarkably, this is just as Waddington predicted 75 years ago, a phenomenon he referred to as canalization [43] and has been observed in crosses between other inbred mouse strains [44,45]. Canalization refers to the likelihood of an organism to proceed towards one developmental outcome, despite variation in the process along the way. This variation can be influenced by among other things the numerous functional genetic variants present in a typical human genome, which may contain thousands of variants that alter gene function [46]. We find that most genetic interactions return trait values to levels seen in control strains, which would act to reduce phenotypic variation among developmental outcomes. This robustness in the face of considerable genetic variation is central to the underlying properties of canalization. These genetic interactions therefore represent a mechanism for storing genetic variation within a population, without reducing individual fitness. This stored genetic variation could then enable populations to more quickly adapt to environmental changes [47].

Finally, the consistently greater effect sizes of main effects relative to average effects suggests that GWAS-type studies consistently underestimate true effect sizes in at least a subset of individuals. Therefore, the key to enabling precision medicine is to identify in which subset of individuals a particular variant has a significant effect. The consideration of epistasis in treatment, although in its infancy, remains a promising avenue for improving clinical treatment regimens, including predicting drug response in tumors [48] and guiding antibiotic drug-resistance [49]. However, true precision medicine will necessitate a more comprehensive understanding of how genetic background, across many loci, affects single variant substitutions.

## Method

### Mice

Chromosome substitution strains (CSS) and control strains were purchased fromThe Jackson Laboratory. These strains include C57BL/6J-Chr3^A/J^/NaJ mice (Stock #004381) (B6.A3), C57BL/6J-Chr4^A/J^/NaJ mice (Stock #004382) (B6.A4), C57BL/6J-Chr5^A/J^/NaJ mice (Stock #004383) (B6.A5), C57BL/6J-Chr6^A/J^/NaJ mice (Stock #004384) (B6.A6), C57BL/6J-Chr8^A/J^/NaJ mice (Stock #004386) (B6.A8), C57BL/6J-Chr10^A/J^/NaJ mice (Stock #004388) (B6.A10), C57BL/6J-Chr14^A/J^/NaJ mice (Stock #004392) (B6.A14), C57BL/6J-Chr17^A/J^/NaJ mice (Stock #004395) (B6.A17) and C57BL/6J (Stock #000664). Mice were maintained by brother-sister matings. All mice used for experiments were obtained from breeder colonies at Case Western Reserve University. Mice were housed in ventilated racks with access to food and water *ad libitum* and maintained at 21°C on a 12-hour light/12-hour dark cycle. All mice were cared for as described under the Guide for the Care and Use of Animals, eighth edition (2011) and all experiments were approved by IACUC and carried out in an AAALAC approved facility. Male mice from strains B6, B6.A4, B6.A5, B6.A10 strains and B6.A8 were bred with female mice from strains B6, B6.A3, B6.A6, B6.A14 and B6.A17 strain. The offspring were weaned at 3 weeks of age. The number of offspring analyzed from each cross is shown in S2 Table for both body weight and plasma glucose, although glucose levels were not measured in one mouse each from the following strains: (B6 × B6.A10)F1, (B6.A14 × B6)F1, (B6.A17 × B6.A10)F1, (B6.A3 × B6.A10)F1, (B6.A6 × B6.A4)F1, (B6.A14 × B6.A5)F1 and (B6.A6 × B6.A5)F1. The mice analyzed from each cross were derived from at least three independent breeding cages. No blinding to the genotypes was undertaken.

### Mouse phenotyping

At 5 weeks of age, mice were fasted 16 hours overnight and body weight was measured. Mice were anesthetized with isofluorane and fasting blood glucose levels were measured via retro orbital bleeds using an OneTouch Ultra2 meter (LifeScan, Milpitas, CA, USA). Mice were subsequently sacrificed by cervical dislocation and the caudate lobe of the liver was collected and immediately placed in RNAlater (Thermo Fisher Scientific, Waltham, MA, USA).

### Trait analysis

To analyze the body weight and fasting plasma glucose data, linear regression was used with a main effects term and a term for each pairwise interaction for the males and females separately. In the glucose data, 5 observations were Winserized by setting a ceiling of 4 median absolute deviations from the median. Any values larger than the ceiling (165 mg/dL) were set to the ceiling. Additionally, interactions where one of the crosses contained less than 5 mice were not analyzed leading to the removal of the (B6.A4 × B6.A3)F1 mice, the female (B6.A8 × B6.A14)F1 and the male (B6.A8 × B6.A3)F1 mice. For each sex, two omnibus tests were performed to see if (1) there were any main effects and (2) there were any interaction effects. Fisher’s method was used to combine the omnibus p-values from males and females[50]. Inverse-variance meta-analysis was used to combine the coefficient estimates from the males and females. To account for potential non-normality, heteroscedasticity and multiple testing, we created 10,000 bootstrap data sets by sampling with replacement from each cross and sex combination. Studentized bootstraps (i.e. using pivotal statistics) were used to create confidence intervals for the coefficients and p-values. Multiple tests were adjusted for by comparing the observed test statistics to the maximum bootstrap test statistic as described[51]. P-values were adjusted for multiple comparisons separately for each trait and separately for the main effects and interactions. The proportion of the genetic variance explained by interactions was estimated as (R_Full_ – R_Additive_)/ R_Full_ where R_Additive_ and R_Full_ are the adjusted coefficients of determination for the model with only main effects and for the full interaction model respectively. The adjusted coefficients of determination are an estimate of the proportion of variation in the trait which is explained by the model. Note that R_Full_ and R_Additive_ share the same denominator (i.e. the total trait variation). Thus, total trait variation cancels out of the quantity (R_Full_ - R_Additive_)/ R_Full_ so that the quantity represents the amount of genetic variation that cannot be explained by main effects only. Using the adjusted version of the coefficient of determination helps account for potential overfitting. Bootstrap confidence intervals of this proportion were calculated.

### Sample preparation for RNA-Seq

Liver tissue stored in RNAlater was homogenized using a Tissumizer Homogenizer (Tekmar, Cincinnati, OH, USA). Total RNA was isolated using the PureLink RNA purification kit (Thermo Fisher Scientific, Waltham, MA, USA). A sequencing library was generated using the TruSeq Stranded Total RNA kit (Illumina, San Diego, CA, USA). RNA samples were sequenced on Illumina HiSeq2500s with single-end 50 base pair reads [52]. Library preparation and RNA sequencing were performed by the CWRU genomics core (Director, Dr. Alex Miron). A total of 7,808,410,316 reads were generated across four flow cells, with an average of 49,735,098 reads per sample. Sequencing quality was assessed by FastQC [53], which identified an average per base quality score of 35.46.

### RNA-Seq data analysis

To maximize statistical power, 20 samples were selected for analysis from the control B6 group, 8 samples were selected from the single CSS groups, and 5 samples were selected from the double CSS groups. A total of 154 control and CSS mice were analyzed, including 20 B6 mice, 63 mice that were heterozygous for one A/J-derived chromosome, and 71 mice that were heterozygous for two different A/J-derived chromosomes. The B6.A4 × B6.A3 and B6.A8 × B6.A3 crosses were poor breeders and thus we did not obtain 5 samples to analyze from these crosses.

Reads were aligned using TopHat2 (2.0.10) [54] to the reference mm10 genome. Because the reference genome is comprised of sequence from strain B6, sequencing reads from a B6-derived chromosome are more accurately mapped than reads from an A/J-derived chromosome [55]. To avoid potential mapping biases, we created an “individualized genome” of the A/J mouse strain using the program Seqnature [55] with variant calls from the Mouse Genomes Project that were downloaded from The Sanger Institute [56]. Reads that were not mapped to the B6 genome were then mapped to the individualized AJ genome with TopHat2. HTSeq-count [57] and the GENCODE vM7 gene annotations[46] were used to count the number of reads for each gene feature. Graphical depictions of the distribution CPM (counts per million) were used to remove 3 outlier samples. Genes where less than 75% of the samples had a count greater than or equal to 15 were considered to be expressed at low levels in liver and were removed leaving 13,289 genes that were considered expressed. To enhance reproducibility and reduce the dependence between the genes, svaseq [59] was used to create 5 surrogate variables that served as covariates in subsequent modeling.

EdgeR [58] was used to fit a model with main effects and pairwise interactions between each chromosome substitution. EdgeR uses a log link function, and thus departure from additivity in EdgeR is departure from a multiplicative model on the gene expression level. The chromosome-chromosome interactions with FDR < 0.05 were divided into the categories synergistic and antagonistic based on the gene expression differences between the double CSS strain and the control strain relative to that predicted by an additive model (S9 Fig.).

To estimate the amount of variation attributable to interaction, we fit an additive model in EdgeR which did not include any interaction terms. We then calculated for each individual and gene the fitted values assuming that the individual’s covariates (i.e. the SVA surrogate variables) were set to 0. We then calculate SS_Full_ as the sum of the mean centered and squared fitted values for the full model including interaction, S_Additive_ was calculated similarly for the additive model. We calculated the amount of the genetic variation explained by the interaction as (SS_Full_ - S_Additive_) / S_Full._ This estimate may be slightly biased upward due to overfitting. However, the mean value for this statistic among the genes with no significant interaction (FDR > 0.5) was 0.25 (1^st^ quartile: 0.20, 3^rd^ quartile: 0.32) (Fig. 5B), which gives an estimate of the upper bound on the possible bias.

### Quantitative PCR (qPCR)

Tissue was homogenized using TissueLyser II (Qiagen, Valencia, CA, USA) and total RNA was isolated using the PureLink RNA purification kit with TRIzol protocol (Thermo Fisher Scientific, Waltham, MA, USA). Total RNA was reverse transcribed using the high capacity cDNA reverse transcription kit (Applied Biosystems, Carlsbad, CA, USA). The sequences for each primer are listed in S13 Table. The qPCR reactions were performed with the power SYBR green PCR Master Mix (Thermo Fisher Scientific, Waltham, MA, USA) and run on a Bio Rad CFX Connect Real Time System (Bio Rad, Hercules, CA, USA). Expression levels were calculated using the ΔΔCt method relative to the *Rplp0* control gene.

### Data availability

The RNA-Seq data is available from GEO under the accession number GSE93591.

## Acknowledgements

This work was supported by the NIH grant DK099533 to D.A.B.. A.C. was supported by a grant from the Sigma Xi Scientific Research Society.

## Author Contributions

D.A.B. and N.M devised the experimental design. D.A.B., A.C., and Y.L. performed the experiments. D.A.B., N.M., S.W., and A.C. analyzed the data. D.A.B, N.M., and A.C. wrote the manuscript. All authors revised the manuscript.

